# Habitat partitioning and spatial segregation at multiple scales promotes year-round coexistence in a guild of forest songbirds

**DOI:** 10.1101/2024.08.26.609673

**Authors:** Alessandro Berlusconi, Giulia Castiglione, Lucas A. Wauters, Alessio Martinoli, Erminio Clerici, Andrea Mologni, Michelangelo Morganti, Adriano Martinoli, Andrea Romano, Diego Rubolini

## Abstract

1. Despite ecological, biological, or genetic similarities, species belonging to the same guild should exhibit some degree of niche differentiation to coexist in the same area. Understanding the mechanisms promoting coexistence among ecologically similar and phylogenetically related species is crucial to improve our understanding of how biodiversity is maintained across broad temporal scales.
2. We examined the mechanisms driving coexistence in a guild of 5 sympatric woodland songbirds (family Paridae) in mixed forests of south-central Europe. We performed interspecific comparisons of habitat and space use considering two different phenological periods (breeding and non-breeding) and two spatial scales (home-range and foraging habitat), as well as spatial segregation of breeding territories.
3. Based on broad-scale habitat preferences, two distinct and seasonally consistent species subgroups were identified within the guild, namely broadleaf and conifer species. During breeding, we showed that all species largely overlapped in their use of different foraging micro-habitats within the tree canopy, even within each subgroup. Yet, we detected significant spatial segregation of breeding territories among species. On the contrary, during the non-breeding period, individuals of different species within mixed flocks foraged on different and complementary sectors of the canopy, thus partitioning foraging habitats.
4. This study highlights that coexistence within the south-central European tit guild across different phenological periods is facilitated by a combination of distinct and complementary mechanisms, including spatial segregation of breeding territories, revealing how sympatric closely-related species coexist through both broad- and fine-scale spatial niche differentiation.

## 1. Introduction

Interspecific competition has long been considered a crucial evolutionary force in regulating population and community structure (Gurevitch *et al*. 2000, Kneitel 2019). Closely related species may show variable intensity of interspecific competition due to shared evolutionary traits and similar ecological requirements (Schluter 2000, Powell *et al*. 2021). Yet, they frequently occur sympatrically, and it is generally assumed that this is associated with ecological niche differentiation (Brown & Wilson 1956, Barabás *et al*. 2016), which ensures long-term coexistence by reducing competition for shared limited resources (MacArthur 1958, Grant 1972). Niche differentiation may be realized through several non-mutually exclusive ways, including differences in dietary preferences, habitat selection or spatial or temporal distribution in feeding/nesting areas (MacArthur 1958, Diamond 1973, Lovette & Hochachka 2006, Bonaccorso *et al*. 2007). Thus, despite sharing the same geographical range, sympatric species may exhibit distinct local distributions by specializing in different (micro)habitat types, i.e. occupying distinct canopy layers or partitioning foraging habitats within a shared forest (MacArthur *et al*. 1972, Demaya *et al*. 2020).

Species within the same guild (i.e. a group of species exploiting the same class of environmental resources, Simberloff & Dayan 1991) may be considered as potential competitors (Simberloff & Dayan 1991, Barabás *et al*. 2016), and their long-term coexistence became viable only if niche differentiation emerges (Martin & Thibault 1996, Bonaccorso *et al*. 2007, Kent & Sherry 2020). The coexistence of ecologically similar species may also be afforded by local spatial segregation, achieved through interspecific territorial behavior (Depino & Areta 2020). According to Drury *et al*. (2020), the evolutionary persistence of interspecific territoriality may act as an alternative to niche differentiation when ecological niches are largely overlapping, especially among phylogenetically related species. Unravelling the ecological dynamics promoting interspecific coexistence is a central goal in ecology and evolution, and it is essential to understand how biodiversity is maintained over broad temporal scales (Chesson 2000). Hence, ecological guilds represent excellent study systems for investigating the mechanisms facilitating the coexistence of similar species across evolutionary times.

Among birds, the Western European tit (family Paridae) includes 6 species that show similar foraging behaviors and diets (Jansson *et al*. 1981, Jansson & Von Brömssen 1981), habitat preferences (Cramp & Perrins 1993), and niche space utilization without habitat separation. This set of Western European species are phylogenetically close and have been coexisting as guild since at least the last post-glacial period (14-16,000 years ago), although it is possible that these species have been in contact for approximately the past ∼5 million years (Tietze & Borthakur 2012, Johansson *et al*. 2013), suggesting that such a long-lasting coexistence may be due to niche differentiation. Despite tits are among the most extensively studied avian taxa (e.g. Savill *et al*. 2010, Dhondt 2012), only a handful of studies have investigated factors promoting coexistence, often focusing on two or at most three species and in specific periods of the annual cycle (e.g. Ekman 1979, Alatalo 1982, Alatalo *et al*. 1986, Hartley 1987, Krams 1996, Dhondt 2012, Maziarz *et al*. 2023).

Here, we aimed at elucidating ecological mechanisms that promotes coexistence in the central European tit guild, consisting of five species belonging to different genera of the Paridae family ( Great Tit *Parus major*, Blue Tit *Cyanistes caeruleus*, Marsh Tit *Poecile palustris,* Crested Tit *Lophophanes cristatus*, Coal Tit *Periparus ater*) throughout the annual cycle. We investigated niche partitioning at various spatial scales to determine if it is achieved through: 1) differential habitat use at the home-range scale during both the breeding (spring) and non-breeding (winter) periods among different species; 2) differential use of sectors in tree canopies by foraging individuals as a proxy for ecological differentiation at the foraging habitat scale in both periods; and 3) spatial segregation of breeding territories among different species during the spring.

## Materials & methods

### Target species and study area

The five species composing the guild object of the study are all commonly found in woodland habitats across central-southern Europe. They are small-sized birds (10-16 cm) which differ mostly in plumage coloration and body size (Gosler & Clement 2007). These species are all mainly associated with forests: the Great Tit, Blue Tit and Marsh Tit typically live in broadleaf forests (Bellamy *et al*. 2000, Hinsley *et al*. 2007) (“broadleaf species” hereafter), while the Crested Tit and the Coal Tit in coniferous forests (Brotons & Herrando 2003, Berlusconi *et al*. 2022) (“conifer species” hereafter). However, the species-specific preferences of habitat use are known to vary according to different ecological and geographical contexts (Blondel 1985, Tremblay *et al*. 2003, Charmantier *et al*. 2017, Pollock *et al*. 2017). These birds have a high metabolic rate and spend most of their time foraging (Lilliendahl *et al*. 1996). Their diet undergoes deep seasonal changes, shifting from insects (Lepidoptera larvae and spiders) in spring, towards seeds, fruits, and buds during the winter (Cramp & Perrins 1993, Gosler & Clement 2007). Tits exhibit complex social behavior: during the breeding period, they form stable and strongly territorial pairs – although extra-pair matings and multiple paternity of nestlings has been demonstrated (Kempenaers *et al*. 1992, Blakey 1994, Dietrich *et al*. 2004) –, while during winter they merge in flocks i.e. both mono-specific and mixed-species groups of individuals (Herrera 1979), displaying erratic movements or forming stable flocks (Ekman 1979, Alatalo *et al*. 1986). The boundaries of breeding territories are bounded by singing males, which defend them until offspring fledge (middle June).

The study was carried out within a 4,828-ha protected forest area named “Pineta Park of Appiano Gentile and Tradate” (Northern Italy, 45°44’ N - 8°56’ E), covered for almost 73% by forests and woodlands (Fig. S1). The area is characterized by intricate fluvioglacial terraces, at altitudes spanning between 243 and 447 m a.s.l.. The woodland is structured in patches of acidophilous forests, primarily consisting of Scots pine *Pinus sylvestris*, and broad-leaved trees (chestnuts *Castanea sativa*, birches *Betula pendula*), as well as oaks (*Quercus petraea, Q. robur*) and hornbeams *Carpinus betulus*. Other patches, especially adjacent to farmlands (Bianchi 2002), are dominated by non-native tree species, mainly black locusts *Robinia pseudoacacia*. This peculiar tapestry of coniferous and broadleaved tree patches fosters a diverse range of resources and habitats. Notably, the study area is among the few localities in Italy where all these five tit species coexist in shared forest patches throughout both the breeding and non-breeding seasons.

### Habitat use during breeding period

#### Home-range scale

Habitat use and differentiation during breeding were studied at the home-range scale by considering the different portions of forest habitats occupied by breeding territories of different species and performing a multinomial analysis for compositional data, testing differences in habitat use among species (Douma & Weedon 2019).

Forest habitats were classified by the dominant species (canopy cover >50%) and by a sub-type (canopy cover >25%) at a 30 m spatial resolution (see Bianchi 2002). Forest habitats were mapped during the 2021 and 2022 vegetative seasons (from June to October). Eventually, we obtained a comprehensive land-use map of the forested areas in the study site. Overall, 43 forest categories were identified at first, subsequently merged into 5 major habitat classes to favor the ecological interpretability of the findings based on both macro-scale structure and typology (see Table S1): acidophilic conifer woods, acidophilic mixed woods, acidophilic broadleaf woods, mesophilic broadleaf woods, alien plant species woods.

We then identified the territories of breeding pairs using territory mapping of unmarked birds (see Bibby *et al*. 2007). We selected 10 territory mapping areas to survey (mean size of 58.6 ± 3.8 ha, Fig. S1), ensuring that they encompassed diverse habitat classes with similar coverage and being characterized by a dense network of paths facilitating monitoring (as suggested by (Bibby *et al*. 2007) each area was surveyed 9 times between March 15^th^ and May 15^th^ 2022 with surveys carried out from sunrise until mid-day and spaced apart by a minimum of 3 days. All individual observations were georeferenced using QGIS version 2.18.28 (QGIS Development Team 2018).

For each species, observations were assigned to putative male territories following the standard methodology outlined by Bibby *et al*. (2007). This involved analyzing clusters of bird observations to determine which points belonged to individual territories. Specifically, clusters with multiple simultaneous registrations of individuals were essential for accurately identifying the points associated with males. Subsequently, bird records were visually assigned to the putative territories of male individuals. At the end of the process, we defined a territory when accomplishing the following criteria: a minimum of 5 points, collected from at least 2 separate surveys and a minimum of 1 behavior indicating the occurrence of a nest (e.g. territorial singing, entering a cavity, food provisioning, etc.) was observed. The minimum convex polygon was used as a method to define the borders of territories (see Hill & Lein 1989, Assandri *et al*. 2018, Juárez Jovel *et al*. 2020), using the “adehabitatHR” R package (Calenge 2011). A total of 456 territories were identified: 164 of Great Tits, 61 of Blue Tits, 118 of Marsh Tits, 61 of Crested Tits and 52 of Coal Tits. Territories were overlaid to the land-use map to determine habitat composition (proportion of the 5 habitat classes described above).

Variability of habitat use of territories among the 5 species was assessed by fitting a Dirichlet model (Maier 2021), with proportional use of habitat classes set as the response variable and the different species set as explanatory variable. This form of regression analysis is suitable for response variables consisting of proportions that sum to one (i.e. compositional data; Douma & Weedon 2019). According to Douma & Weedon (2019), we replaced all observed zeros with a small number (equal to the smallest non-zero observation for each habitat class) prior to fitting the model. This analysis was carried out using the R package “DirichletReg” (Maier 2021). All analyses were performed using R version 4.2.0 (R Core Team 2023).

#### Foraging habitat scale

Habitat use at the foraging habitat scale was assessed by comparing both preferences for tree-type and selected sectors of the tree canopy among foraging individuals of different species (Lens *et al*. 1994, Hinsley *et al*. 2007). Foraging birds on tree canopies were recorded using instantaneous scan sampling (Altmann 1974). To reduce statistical dependence of observations, we only recorded one foraging position per individual (see Lens *et al*. 1994). The tree-type on which they foraged was classified as “broadleaf” or “conifer”. We then assigned the vertical location within canopy along a three-level factor: top third; middle; bottom. Analogously, horizontal location within the canopy was defined with a further three-level factor: inner, middle or outer. Breeding period data were collected during 2021 and 2023 (March 15^th^ – May 15^th^) by collecting observations along the same paths used for territory mapping. Surveys were carried out in the morning, from sunrise to mid-day. We collected a total of 201 observations concerning Great Tits, 152 for Blue Tits, 167 for Marsh Tits, 138 for Crested Tits and 76 for Coal Tits.

The distribution (i.e. count) of observations for each species in broadleaf or coniferous trees and then in each of the different sectors of the canopy were used as the dependent variables to indicate foraging habitat preference. The species were assigned to 2 subgroups (“broadleaf” and “conifer species”) using Pearson’s χ^2^ test. Subsequently, within each group, we compared the distribution among species in the sectors of the canopy used during foraging. The Cochran-Mantel-Haenzsel (CMH) test, designed to explore the significancy of unevenly distribution in three-ways contingency tables, was applied to evaluate distinctions in species canopy distribution, with *post hoc* analyses performed using Fisher’s exact test.

#### Spatial segregation of breeding territories

We quantified interspecific spatial segregation among breeding territories at the species level. Initially, for each mapping area, we calculated the total surface area covered by breeding territories of each species. Subsequently, we calculated the percentage of overlap (% overlap hereafter) for each species relative to other ones within “broadleaf” and “conifer species” subgroups separately, using the “sf” package for R (Pebesma 2018). For each species, we determined the percentage of its territory covered by those of other species. We first tested if % overlap did depend on density of territories across mapping areas, by correlating the average % overlap to the territory density (territories/ha of suitable area) of each subgroup in each mapping area (see Appendix 1 in Supporting Information for details). Then, we tested whether the observed overlap values were different from those expected by chance at each density of breeding territories, thus in the absence of spatial segregation. To this end, we compared the observed values of overlap against simulated ones within each mapping area. Briefly, for each mapping area the observed territories for each species were randomly repositioned within the suitable available habitat (i.e. territories of “broadleaf species” were relocated over all broad-leaved woods and acidophilic mixed woods, while those of “conifer species” over acidophilic conifer and acidophilic mixed woods). We then calculated the % overlap for each species compared to others within each subgroup. This process was reiterated 100 times in each mapping area (see Appendix 2 in Supporting Information for details). Finally, observed and randomized territory overlaps were compared through a Wilcoxon signed-rank tests for paired data, for each mapping area (N = 10 “broadleaf”; 8 “conifer species”)

### Habitat use during the non-breeding period

#### Home-range scale

Because these species do not exhibit territoriality during the non-breeding period, the whole study area was considered as the home-range for all species. Data were collected during the 2020-2021 and 2021-2022 winter (December 1^st^ – February 10^th^). As for the breeding period, we relied on instantaneous scan sampling to collect observations of foraging individuals, recording a single location per individual. “Locations” were defined as any location where an individual was observed foraging, preening, or calling. Observations were conducted along 60 transects (5-8 km long), mostly along trails and roads, across the whole study area. Transects were surveyed in the morning (after sunrise). Random opportunistic observations were also considered. A total of 1,208 locations were recorded with this method, respectively belonging to: Great Tit (N=364), Blue Tit (N=172), Marsh Tit (N=235), Crested Tit (N=232) and Coal Tit (N=205).

To assess habitat preferences, we computed Manly’s selection ratios for each species in each habitat class within the study area, based on occurrence locations (Manly *et al*. 2004). Selection ratios were calculated accordingly to a use vs. availability design, where the selection ratio (i.e. Manly’s selection ratio) is defined as the ratio between use and availability measures. We used as “availability” of a given habitat class and its proportional availability in the whole study area. Habitat “use” was calculated as proportion of locations assigned to a given habitat class with respect to the total number of locations. Selection ratios along with their 95% confidence intervals (95% CI) were estimated using Koopman’s score method (Aho & Bowyer 2015) using the “asbio” R package (Aho 2014).

#### Foraging habitat scale

Foraging habitat use during the non-breeding period was evaluated as for breeding. Data were collected during 2020-2021 and 2021-2022 (December 1^st^ – February 10^th^), obtaining a total 705 observations: Great Tits (N=188), Blue Tit (N=109), Marsh Tit (N=162), Crested Tit (N=119), Coal Tit (N=127). We also compared the differences in the distribution in tree canopy during foraging between in-flock individuals and singletons within each species during the non-breeding period with CMH tests.

## Results

### Habitat use during breeding period

#### Home-range scale

During breeding, territories of Great Tits, Blue Tits and Marsh Tits were mainly composed of broadleaf woods (Fig. 1), especially mesophilic and secondarily by alien plant species woods. In contrast, Crested Tits and Coal Tits preferred habitats dominated by conifers (Fig. 1). Habitat use significantly differed among species (Table S2). Yet, no statistically significant differences were observed among Great Tits, Blue Tits, and Marsh Tits (the “broadleaf species”), nor between Coal Tits and Crested Tits (the “conifer species”) (Table S2). Hence, within conifer and broadleaf species groups, all species shared similar proportion of habitat classes, indicating overlap in habitat use.

**Figure 1.**
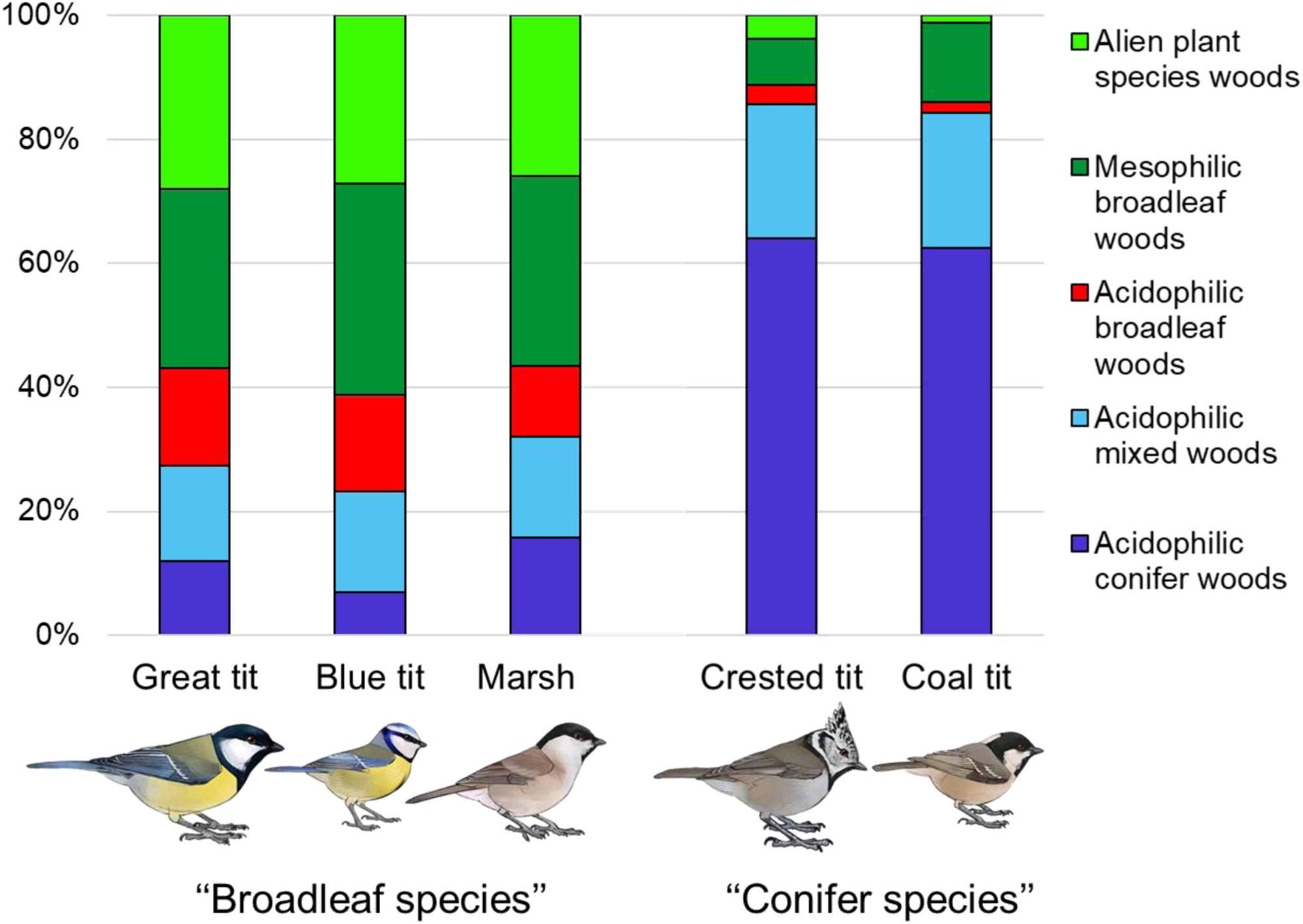
Broad-scale habitat use during breeding of 5 European tit species, characterized by the % habitat composition within breeding territories. The species are clearly subdivided into two distinct subgroups: broadleaf species (Great, Blue and Marsh Tits) and conifer species (Crested and Coal Tits).

#### Foraging habitat scale

Great Tits, Blue Tits, and Marsh Tits showed a strong preference for broadleaf trees, whereas Crested Tits and Coal Tits preferred conifers (Table 1). These findings match those obtained at the home range scale.

**Table 1.** Broad-scale habitat preferences for broadleaf vs. conifer trees as foraging substrate. A strong preference and seasonally coherent preference for broadleaf trees emerged for Great, Blue, and Marsh Tits, while Crested and Coal Tits preferred conifers.

| Species | Ratio of broadleaf<br>vs. conifer | $\chi^2$ | df | *P* |
| --- | --- | --- | --- | --- |
| *a) Breeding period* |  |  |  |  |
| Great Tit | 9.00 | 128.96 | 1 | < 0.001 |
| Blue Tit | 24.33 | 128.95 | 1 | < 0.001 |
| Marsh Tit | 9.44 | 86.26 | 1 | < 0.001 |
| Crested Tit | 0.29 | 41.85 | 1 | < 0.001 |
| Coal Tit | 0.55 | 6.37 | 1 | 0.012 |
| *b) Non-breeding period* |  |  |  |  |
| Great Tit | 3.33 | 41.46 | 1 | < 0.001 |
| Blue Tit | 12.57 | 76.05 | 1 | < 0.001 |
| Marsh Tit | 2.11 | 23.73 | 1 | < 0.001 |
| Crested Tit | 0.05 | 80.42 | 1 | < 0.001 |
| Coal Tit | 0.15 | 41.81 | 1 | < 0.001 |

Regarding canopy utilization, no differences emerged between the two conifer species (Table 2, Fig. 2), whereas broadleaf species showed significant distribution patterns, unevenly within the canopy (Table 1, Fig. 2). Specifically, *post hoc* tests clarified that all the species avoiding the outer sector of the canopy, Great Tits used evenly the middle and inner sectors, and Blue and Marsh Tits being more frequent in the middle sector (Fig. 2).

**Figure 2.**
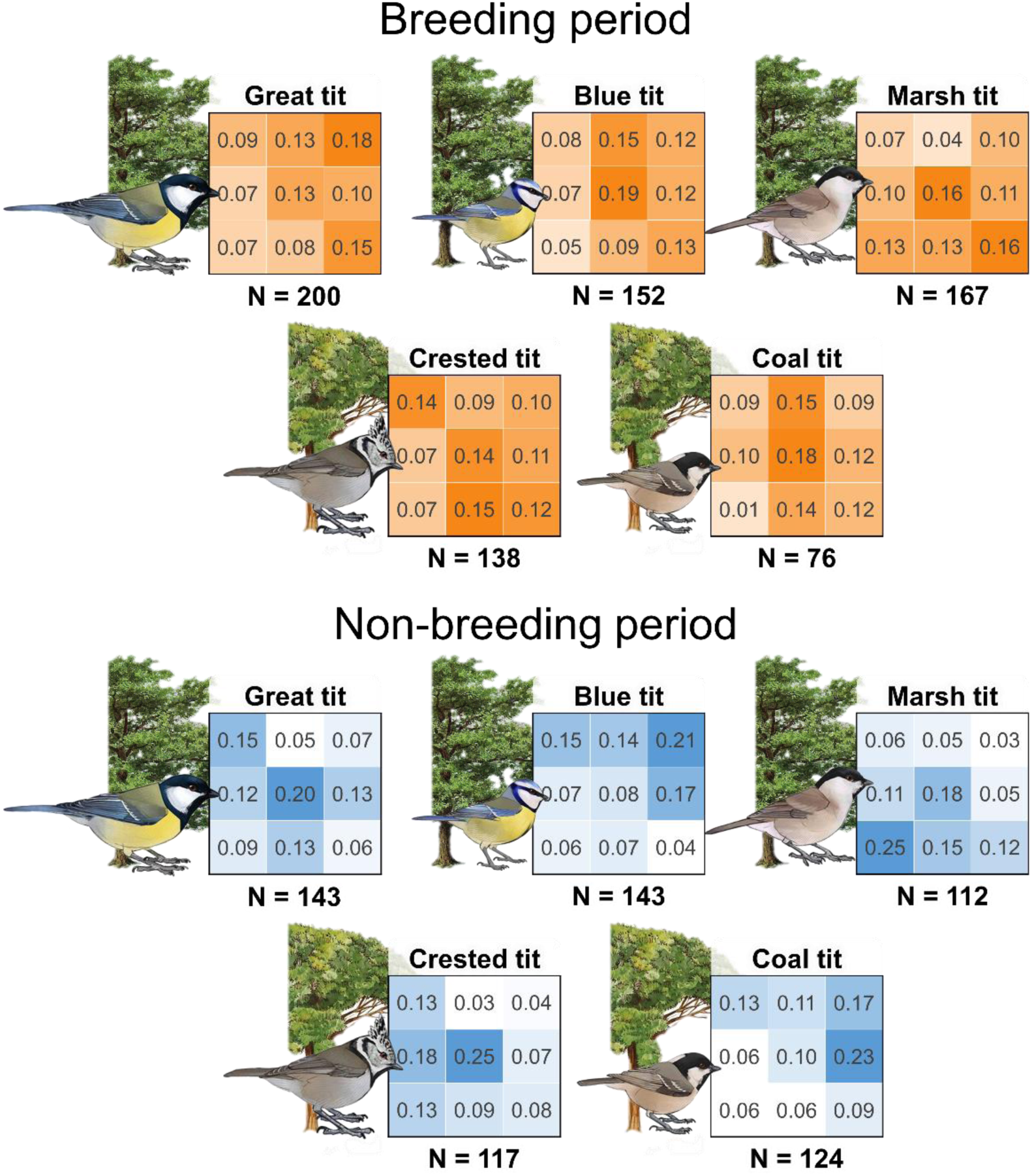
Fine-scale partitioning of foraging microhabitat within the tree canopy, expressed as the frequency of foraging individuals across different sectors of the tree canopy during the breeding (upper panel) and non-breeding (lower panel) periods for broadleaf (upper sub-panel of each panel) and conifer species (lower sub-panel). The tree canopy was divided into 9 distinct sectors, by combining vertical – “top third”, “middle third” and “bottom third” – and horizontal sectors – “inner third”, “middle third” and “outer third”. Numbers represent frequency of foraging individuals per each canopy sector. Total number of observations is presented below each table. In the breeding period, all species overlapped in their use of the entire canopy even within subgroups; however non-breeding species fed on different tree-types according to their subgroups, using complementary sectors of the canopy, effectively partitioning space use and promoting coexistence.

**Table 2.**
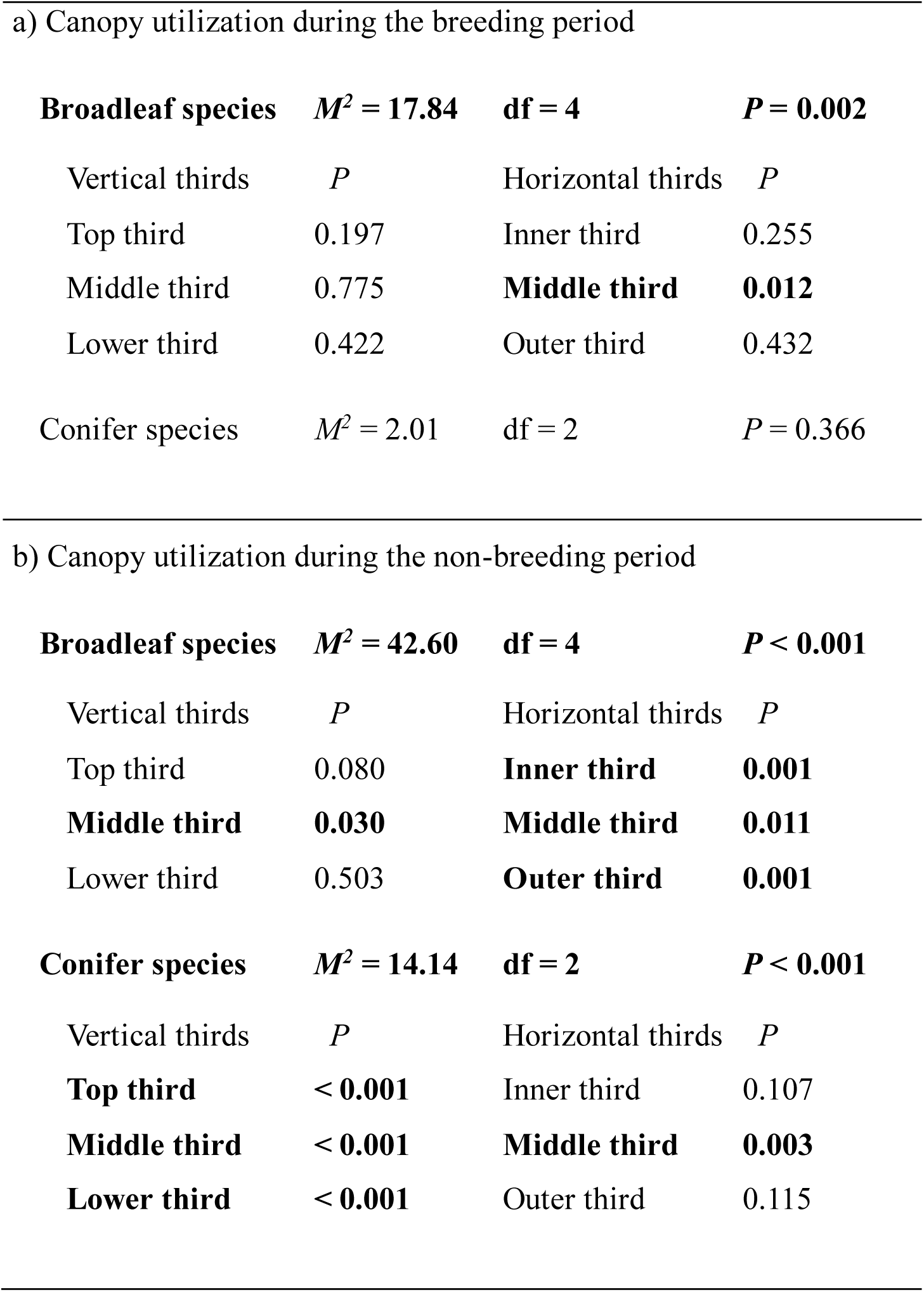
Fine-scale habitat preferences (frequency of canopy utilization) during a) breeding and b) non-breeding period (see Fig. 3 for frequency data). Cochran-Mantel-Haenszel (CMH) tests were performed separately within each subgroup (“broadleaf” and “conifer species”). If the CMH test is significant, *post hoc* analyses were performed using Fisher’s exact test. Results of *post hoc* tests are shown to identify the sectors of the canopy where the distribution of species differed significantly. Bolded text denotes statistically significant variables (*P* < 0.05).

|  |  |  |  |
| --- | --- | --- | --- |
| a) Canopy utilization during the breeding period |  |  |  |
| **Broadleaf species** | **$M^2 = 17.84$** | **df = 4** | **$P = 0.002$** |
| Vertical thirds | $P$ | Horizontal thirds | $P$ |
| Top third | 0.197 | Inner third | 0.255 |
| Middle third | 0.775 | **Middle third** | **0.012** |
| Lower third | 0.422 | Outer third | 0.432 |
| Conifer species | $M^2 = 2.01$ | df = 2 | $P = 0.366$ |
| b) Canopy utilization during the non-breeding period |  |  |  |
| **Broadleaf species** | **$M^2 = 42.60$** | **df = 4** | **$P < 0.001$** |
| Vertical thirds | $P$ | Horizontal thirds | $P$ |
| Top third | 0.080 | **Inner third** | **0.001** |
| **Middle third** | **0.030** | **Middle third** | **0.011** |
| Lower third | 0.503 | **Outer third** | **0.001** |
| **Conifer species** | **$M^2 = 14.14$** | **df = 2** | **$P < 0.001$** |
| Vertical thirds | $P$ | Horizontal thirds | $P$ |
| **Top third** | **< 0.001** | Inner third | 0.107 |
| **Middle third** | **< 0.001** | **Middle third** | **0.003** |
| **Lower third** | **< 0.001** | Outer third | 0.115 |

#### Breeding density and spatial segregation of breeding territories

Overall, Great Tits showed the highest density of breeding territories (mean ± SE: 0.35 ± 0.06 territories/ha), followed by Marsh Tits (0.23 ± 0.03), Crested Tits (0.17 ± 0.02), Blue Tits (0.12 ± 0.01) and Coal Tits (0.12 ± 0.02). The density of breeding territories was higher among “broadleaf” (0.23 ± 0.03 territories/ha) than “conifer species” (0.15 ± 0.02 territories/ha; Mann-Whitney test, *W* = 165, *P* = 0.048, n = 10 areas). The % overlap of territories among “broadleaf species” (8.82 ± 0.85 %) resulted to be significantly larger than those among the “conifer species” (4.31 ± 1.10 %; Mann-Whitney test, *W* = 1, *P* < 0.001, n = 10 areas, see Table S3). The % overlap of territories across mapping areas was not significantly associated with territory density neither for “broadleaf” nor for “conifer species” (see Appendix 1).

The observed overlap of territories was significantly lower than expected by chance for all species pair comparisons (all *P* < 0.04; Table S4, Fig. S5), suggesting consistent spatial segregation among all species within each of the two subgroups.

### Habitat use during the non-breeding period

#### Home-range scale

Great Tits, Blue Tits, and Marsh Tits exhibited a significant overlap in their habitat preferences, distinct from those of Crested Tits and Coal Tits, which also displayed overlapping selection ratios (Fig. 3). However, there was an exception in the case of acidophilic mixed woods, since this habitat class was selected similarly by the entire guild (Fig. 3) (see also Table S5 in Supporting Information).

**Figure 3.**
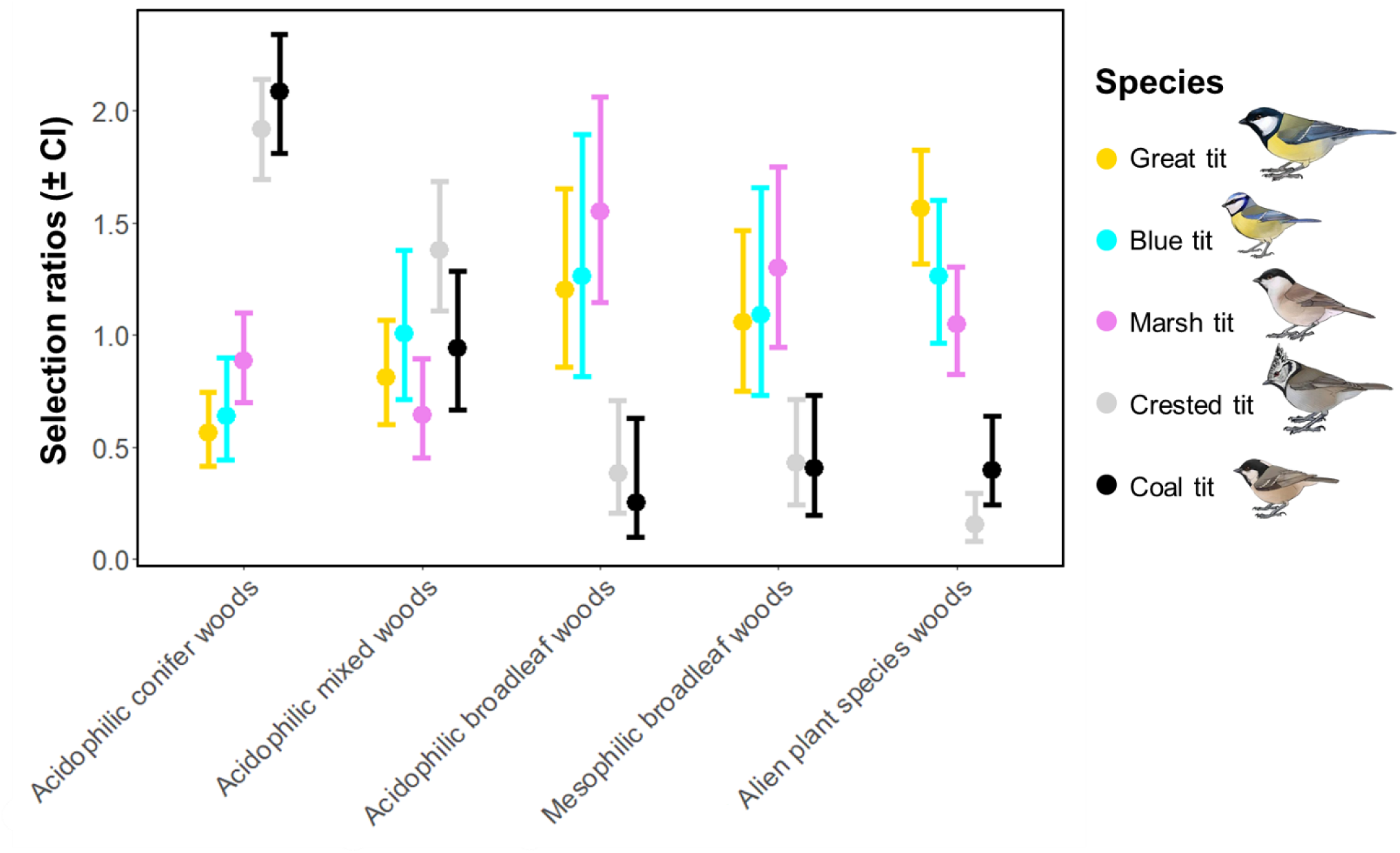
Home-range scale habitat selection during the non-breeding period (Manly’s selection ratios with 95% CI). Both broadleaf and coniferous species exhibited overlapping habitat preferences within their respective groups, while significant differences in habitat preferences were observed between these two groups for most habitat types.

#### Foraging habitat scale

Similar to the breeding period, Great, Blue, and Marsh Tits showed a strong preference for broadleaf trees, whereas Crested and Coal Tits preferred conifers (Table 1). “Broadleaf species” and “conifer species” both exhibited a non-uniform use of the tree canopy (*M^2^* = 42.60, df = 4, *P* < 0.001 and *M^2^*= 14.14, df = 2, *P* < 0.001, respectively) (Table 1, Fig. 2). Within their respective groups, each species displayed a significant preference for distinct canopy sections relative to the other species, suggesting complementary utilization patterns. Notably, all the species exhibited selective preferences for specific sectors: Great Tits preferred central sectors of broadleaf trees, Blue Tits the uppers and outers and Marsh Tits the inners and lowers, in contrast, Crested Tits favored inner and central sectors of conifer trees and Coal Tits the uppers and outers (Table 1, Fig. 2).

Moreover, solitary foraging individuals of each species did not differ in their tree canopy use compared to those observed in flocks (*P* > 0.05 for all comparisons) (see Appendix 3 and Table S6).

## Discussion

Our study of year-round habitat and space partitioning in a guild of 5 species of European woodland tits revealed different mechanisms of small-scale niche differentiation, some of which varied between breeding and non-breeding periods. Besides confirming the traditional distinction between the broadleaf and conifer species, we found inter-seasonal consistency in preferred habitats at the home-range scale within each species’ subgroup. At a finer scale, during the non-breeding period, different tit species foraged in distinct and complementary sectors of tree canopies. In contrast, during the breeding period, all species exploited the entire canopy, thus showing a large spatial niche overlap. In this case, coexistence seemed to be facilitated by limiting the interspecific spatial overlap of breeding territories. Therefore, different mechanisms operate in different moments of the life-cycle to maintain niche segregation: this are territoriality in spring and unevenly distribution among canopy in winter. Our study is the first comprehensive demonstration of these phenomena in the guild of Paridae of Western Europe and one of the rare cases on which this has been demonstrated on vertebrates at all.

The differentiation between “broadleaf” and “conifer species” constitutes the main niche clusterization within the guild, with Great, Blue, and Marsh Tits exhibiting a consistent preference for mesophilic and acidophilic broadleaf woodlands over coniferous ones during the entire circannual cycle (Bellamy *et al*. 2000, Naef-Daenzer *et al*. 2000). Conversely, Crested Tits and Coal Tits preferred habitats dominated by conifers all over the year-cycle, including both pure and mixed coniferous woodlands (Ekman 1979, Berlusconi *et al*. 2022). This clear habitat distinction is mirrored by evolutionary divergence in bill morphology, which determines preferences for broadleaf or coniferous trees. This divergence arises from the need to forage among pine needles or the leaves of deciduous plants (Suhonen *et al*. 1994): species inhabiting coniferous habitats possess more slender bills compared to those in deciduous habitats, a pattern observed in other bird species as well (Snow 1954). Notably, bill morphology as a highly sensitive ecological trait to exploitative competitive pressure (Brown & Wilson 1956, Schoener 1965), changing rapidly and leading to niche partitioning.

During winter, tits were frequently observed foraging in mixed-species flocks (Ekman 1979, Alatalo *et al*. 1986). However, within the subgroups, each species demonstrated a preference for distinct sectors of the tree canopy, effectively partitioning space use and favoring coexistence. These findings align with previous studies (Morse 1978, Hartley 1987, Hinsley *et al*. 2007), which suggested that canopy sector selection is driven by subordinate-dominant dynamics, where dominant and larger species occupy the most predator-protected sites (i.e. innermost sectors of the canopy) (Krams 1996). Yet, in our study system, solitary foraging individuals did not change their foraging behavior compared to those observed in flocks. Therefore, distinct canopy use by different species is likely attributed to their divergent foraging strategies, reflecting fine-scale niche partitioning. An alternative interpretation, proposed by Suhonen *et al*. (1994), associates the canopy position with the distinct body size and morphology of the various tit species. Specifically, smaller species, such as the Blue Tits and Coal Tits, are predicted to forage in the outermost parts of the tree canopy, on thinner branches, whereas larger species such as the Great Tit and Crested Tit feed in the innermost sectors, targeting larger branches, as we observed. The different exploited canopy sectors during winter likely facilitate the co-occurrence of the species by minimizing trophic competition (Diamond 1973), and may be crucial in this resource-limited period, as the species spend most of their time foraging (Gibb 1954). The results from the breeding period, however, depict a distinct scenario. Within each subgroup, the species did not exhibit interspecific differences in foraging behavior, utilizing the entire tree canopy without spatial partitioning. The large overlap may be related to the greater food requirements during the nesting period, when parents must find sufficient food both for self-maintenance and offspring provisioning (Minot 1981). Consequently, efficient exploitation of the entire tree canopy may become essential for securing an adequate amount of food (Alatalo 1982, Szekely 1985).

Therefore, the key to coexistence during breeding is clearly to be found in the spatial segregation of territories among (as well as within-) species. We indeed observed a remarkably low interspecific overlap in territory boundaries among all species 8.82% for “broadleaf” and 4.31% for “conifer species”. Previous studies of other passerine guilds have documented relatively higher interspecific territory overlap, typically ranging from 30% to 70% in Northern American chickadees (Hill & Lein 1989) and 50% in Mediterranean warbler (Pons *et al*. 2008). On the other hand, it is noteworthy that the observed spatial segregation likely arises from the low density of all species in the study area, however occurring with a significantly (and markedly) lower percentage of overlap compared to the simulation. However, we are aware that our estimates of territory overlap must be considered with caution, since we did not have marked birds, and thus could not recognize territorial individuals. Nevertheless, we employed rigorous sampling methods and collected a substantial number of observations across a broad temporal and spatial range, yet providing compelling evidence that the study outcomes are robustly representative across the study area.

Some authors have proposed that limited territory overlap is affected by different habitat preferences at the home-range scale (Martin & Thibault 1996). Yet, observed habitat preferences were remarkably similar within each subgroup at both the home-range and foraging habitat scales. We cannot rule out that latent not-studied micro-habitat characteristics could have played a role in niche partitioning during the breeding period. Otherwise, the utilization of food resources by individuals of other species within the territory of a given species could induce competition for those resources (Minot 1981), potentially triggering interspecific aggression and territoriality (Jankowski *et al*. 2010), resulting in spatially segregated territories among species. Interspecific territoriality has not yet been documented in European tits, and this behavioral strategy is often facultative and dependent on local context and breeding densities (Drury *et al*. 2020), potentially arising in local communities according to the level of both intra- and interspecific competition (MacArthur 1958, Drury *et al*. 2020).

Intra-guild niche partitioning generally occurs among co-evolved species, through evolutionary or phenotypically plastic responses. Within shared habitats, coexisting species may either shift ecological requirements through the action of natural selection, progressively acquiring so-called divergent post-competitive realized niches (Elton & Miller 1954), or adjust their territorial behavior in the presence of competitors through phenotypically plastic responses which limit competition (Pfennig & Murphy 2002, Grether *et al*. 2009). Our multi-level approach sheds light on the ecological mechanisms allowing intra-guild coexistence within the guild of European tits, where coexistence was fostered by a combination of several concurrent and non-mutually exclusive mechanisms, including broad-scale ecological differentiation, plastic behavioral responses to seasonal and environmental variations, and fine-scale resource partitioning. Same mechanisms may be acting in other guilds, highlighting the importance of multifaceted strategies in facilitating species coexistence. Understanding these dynamics is crucial for conservation efforts, as it emphasizes the need to preserve habitat heterogeneity and ecological complexity to maintain biodiversity.

## Acknowledgments

We thank Pineta Park of Appiano Gentile and Tradate (ATE Insubria-Olona), specifically G. Pozzi and M. Clerici, who made the project possible. Thanks to A. Patta, coordinator of the Park “Fauna Group”, and all the other Volunteer Ecological Guards who collaborated in the data collection: C. Bottinelli, A. Barozzi, E. Carugo, F. Speroni, F. Calasso, F. Sabaino, F. Benaglio, G. Berlusconi, I. Botta, L. Tombolato, M. Pagani, M. Maldifassi and S. Colaone. Thanks also to the volunteers of Insubric Group of Ornithology: A. Stocchetti, A. Zarbo, A. Castiglioni, D. Perolini, D. dall’Osto, N. Larroux and R. Pigni. This study was partly supported by the PRIN 2022 funding scheme (project WARMBREED, 2022CWMRNH, CUP Master: G53D23002610006; grant number 2022CWMRNH to A.R.). The authors acknowledge the support of NBFC to CNR, funded by the Italian Ministry of University and Research, PNRR, Missione 4 Componente 2, “Dalla ricerca all’impresa”, Investimento 1.4, Project CN00000033. This research and the experiments comply with the current laws of the country in which they were performed (Italy).

## Author contributions

A.B., L.A.W., Ad.M., Al.M., A.R., D.R. conceived the ideas and designed methodology; A.B., G.C.,

E.C., A.M. collected the data; A.B., G.C., Al.M., An.M., A.R., D.R. analyzed the data; A.B., G.C., L.A.W., M.M., A.R., D.R. led the writing of the manuscript. All authors contributed critically to the drafts and gave final approval for publication.

## Data availability statement

The datasets used will be made available through the Dataverse repository upon acceptance.

## Supplementary Online Material

Additional supporting information may be found online in the Supplementary Online Material section.

