## Supplementary_online_material for "Habitat partitioning and spatial segregation at multiple scales promotes year-round coexistence in a guild of forest songbirds"

### List of contents:

**Figure S1:** Map of the 10 mapping areas identified within the study area. Pag. 3

**Table S1:** Reclassification of land-use categories in new habitat classes. Pag. 4

**Table S2:** Summary statistics of the Dirichlet regression model. Pag. 6

**Table S3:** Proportion of boundary overlap for each species pair within each mapping area. Pag. 8

**Appendix 1:** Description of analysis of influence of territory density on species territory overlap. Pag. 10

**Appendix 2:** Detailed description of simulation of breeding territories. Pag. 11

**Figure S2:** Example of identification of an area of simulation. Pag. 14

**Figure S3:** Example of relocation of breeding territories. Pag. 15

**Figure S4:** Example of territory overlap within observed territories and simulations. Pag. 16

**Table S4:** Wilcoxon test for differences in the % overlap of observed vs. relocated territories. Pag. 17

**Figure S5:** Comparison of % of overlap between observed and relocated breeding territories. Pag. 18

**Table S5:** Manly’s selection ratios and the 95% CI for habitat selection at home-range scale. Pag. 20

**Appendix 3:** Description of analysis of canopy distribution during solitary vs. flocking foraging Pag. 21

**Table S6:** CMH test for differences in the distribution of solitary vs. flocking foraging. Pag. 22

**References** Pag. 23

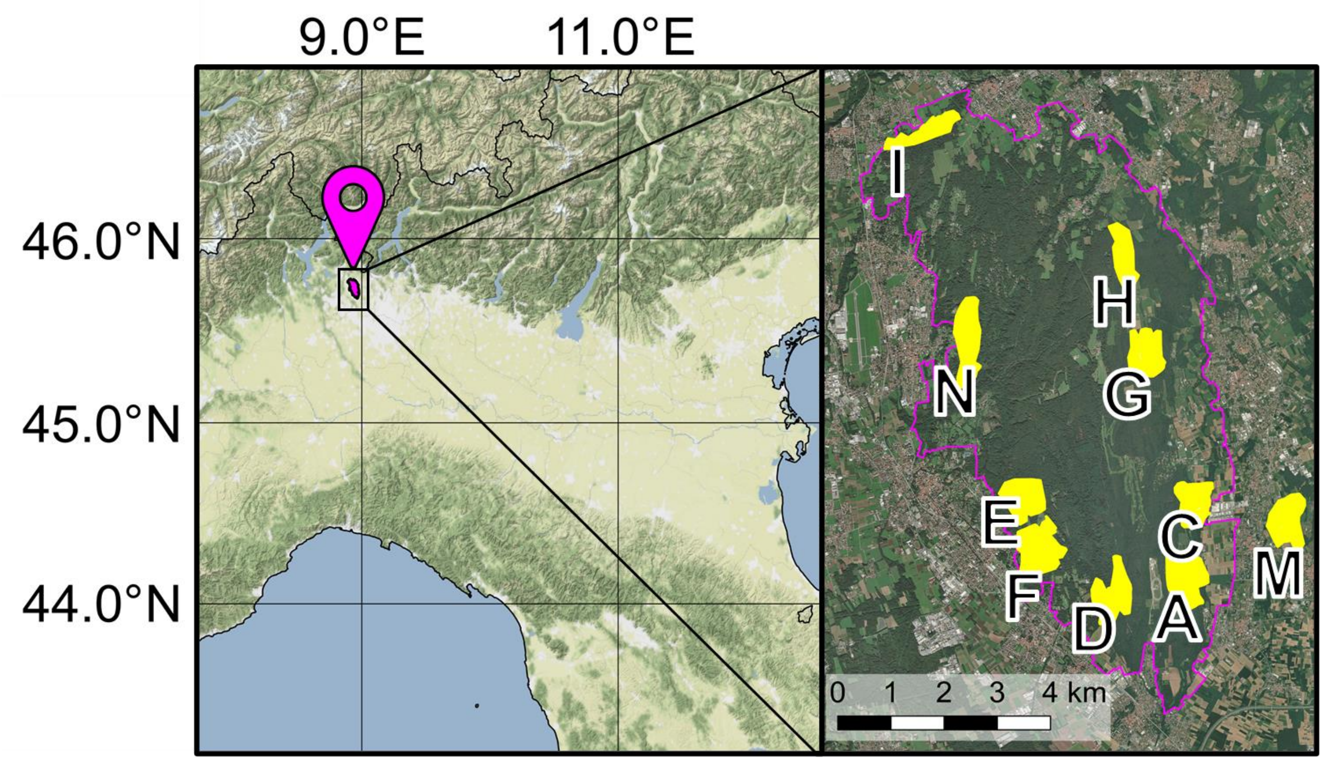


**Figure S1.** Location of the study area in northern Italy (left), with a satellite-view of the Pineta Regional Park of Appiano Gentile and Tradate with the 10 territory mapping areas (right).

**Table S1.** Reclassification of forest categories in new habitat classes used for habitat partitioning at the home range scale.

Reclassification of woodland types

| **Dominant species** | **Sub-type** | **Habitat class** |
|---|---|---|
| *Alnus glutinosa* | *Alnus glutinosa* | Acidophilic broadleaves woods |
| *Acer montanus* | *Acer montanus* | Mesophilic broadleaves woods |
| *Carpinus betulus* | *Carpinus betulus* | Mesophilic broadleaves woods |
| *Castanea sativa* | *Alnus glutinosa* | Acidophilic broadleaves woods |
| *Castanea sativa* | *Carpinus betulus* | Acidophilic broadleaves woods |
| *Castanea sativa* | *Castanea sativa* | Acidophilic broadleaves woods |
| *Castanea sativa* | *Pinus sylvestris* | Acidophilic mixed woods |
| *Castanea sativa* | *Quercus rubra* | Acidophilic broadleaves woods |
| *Castanea sativa* | *Quercus robur* | Acidophilic broadleaves woods |
| *Castanea sativa* | *Robinia pseudoacacia* | Acidophilic broadleaves woods |
| *Pinus sylvestris* | *Carpinus betulus* | Acidophilic conifer woods |
| *Pinus sylvestris* | *Castanea sativa* | Acidophilic conifer woods |
| *Pinus sylvestris* | *Pinus sylvestris* | Acidophilic conifer woods |
| *Pinus sylvestris* | *Quercus rubra* | Acidophilic conifer woods |
| *Pinus sylvestris* | *Quercus robur* | Acidophilic conifer woods |
| *Pinus sylvestris* | *Robinia pseudoacacia* | Acidophilic conifer woods |
| *Prunus serotina* | *Prunus serotina* | Alien plant species woods |
| *Prunus serotina* | *Robinia pseudoacacia* | Alien plant species woods |
| *Quercus rubra* | *Castanea sativa* | Alien plant species woods |
| *Quercus rubra* | *Pinus sylvestris* | Acidophilic mixed woods |
| *Quercus rubra* | *Quercus rubra* | Alien plant species woods |
| *Quercus rubra* | *Quercus robur* | Alien plant species woods |
| *Quercus rubra* | *Robinia pseudoacacia* | Alien plant species woods |
| *Quercus robur* | *Robinia pseudoacacia* | Mesophilic broadleaves woods |
| *Quercus robur* | *Alnus glutinosa* | Mesophilic broadleaves woods |
| *Quercus robur* | *Carpinus betulus* | Mesophilic broadleaves woods |
| *Quercus robur* | *Castanea sativa* | Mesophilic broadleaves woods |

| | | |
|---|---|---|
| *Quercus robur* | *Pinus sylvestris* | Acidophilic mixed woods |
| *Quercus robur* | *Quercus rubra* | Mesophilic broadleaves woods |
| *Quercus robur* | *Quercus robur* | Mesophilic broadleaves woods |
| *Quercus robur* | *Robinia pseudoacacia* | Mesophilic broadleaves woods |
| *Robinia pseudoacacia* | *Alnus glutinosa* | Alien plant species woods |
| *Robinia pseudoacacia* | *Acer montanus* | Alien plant species woods |
| *Robinia pseudoacacia* | *Carpinus betulus* | Alien plant species woods |
| *Robinia pseudoacacia* | *Castanea sativa* | Alien plant species woods |
| *Robinia pseudoacacia* | *Prunus serotina* | Alien plant species woods |
| *Robinia pseudoacacia* | *Pinus sylvestris* | Acidophilic mixed woods |
| *Robinia pseudoacacia* | *Quercus rubra* | Alien plant species woods |
| *Robinia pseudoacacia* | *Quercus robur* | Alien plant species woods |
| *Robinia pseudoacacia* | *Robinia pseudoacacia* | Alien plant species woods |

Reclassification of other land-use types in the study area

| **Other land-use categories** | **New habitat class** |
|---|---|
| Pioneer vegetation | Acidophilic broadleaves woods |
| Riparian vegetation | Acidophilic broadleaves woods |
| Coniferous plantations | Acidophilic conifer woods |

**Table S2.** Summary statistics of the Dirichlet regression model, showing the multiple pairwise comparison among species for each habitat class. The estimated coefficients are reported on a multinomial logit scale. No statistically significant differences were observed in habitat use among Great Tits, Blue Tits, and Marsh Tits (i.e. broadleaf species), nor between Coal Tits and Crested Tits (i.e. conifer species).

| **Habitat class** | **Species pair** | **Estimate ± S.E.** | ***Z*** | ***P*** |
|---|---|---|---|---|
| Acidophilic conifer woods | Great Tit vs Marsh Tit | -0.05 ± 0.12 | -0.42 | 0.678 |
| Acidophilic conifer woods | Great Tit vs Blue Tit | 0.06 ± 0.15 | 0.40 | 0.689 |
| Acidophilic conifer woods | Great Tit vs Coal Tit | -1.20 ± 0.17 | -6.90 | < 0.001 |
| Acidophilic conifer woods | Great Tit vs Crested Tit | -0.91 ± 0.16 | -5.81 | < 0.001 |
| Acidophilic conifer woods | Marsh Tit vs Blue Tit | 0.11 ± 0.16 | 0.70 | 0.483 |
| Acidophilic conifer woods | Marsh Tit vs Crested Tit | -0.86 ± 0.17 | 0.17 | < 0.001 |
| Acidophilic conifer woods | Marsh Tit vs Coal Tit | -1.15 ± 0.18 | -6.32 | < 0.001 |
| Acidophilic conifer woods | Blue Tit vs Crested Tit | -0.97 ± 0.19 | -5.22 | < 0.001 |
| Acidophilic conifer woods | Blue Tit vs Coal Tit | -1.26 ± 0.20 | -6.28 | < 0.001 |
| Acidophilic conifer woods | Crested Tit vs Coal Tit | -0.29 ± 0.21 | -1.42 | 0.156 |
| Mixed conifer woods | Great Tit vs Marsh Tit | -0.08 ± 0.12 | -0.61 | 0.541 |
| Mixed conifer woods | Great Tit vs Blue Tit | -0.03 ± 0.15 | -0.19 | 0.846 |
| Mixed conifer woods | Great Tit vs Coal Tit | -0.32 ± 0.17 | -1.90 | 0.058 |
| Mixed conifer woods | Great Tit vs Crested Tit | -0.22 ± 0.15 | -1.43 | 0.152 |
| Mixed conifer woods | Marsh Tit vs Blue Tit | 0.05 ± 0.16 | 0.29 | 0.770 |
| Mixed conifer woods | Marsh Tit vs Crested Tit | -0.14 ± 0.16 | -0.88 | 0.378 |
| Mixed conifer woods | Marsh Tit vs Coal Tit | -0.24 ± 0.18 | -1.38 | 0.169 |
| Mixed conifer woods | Blue Tit vs Crested Tit | -0.19 ± 0.18 | -1.03 | 0.301 |
| Mixed conifer woods | Blue Tit vs Coal Tit | -0.29 ± 0.20 | -1.48 | 0.140 |
| Mixed conifer woods | Crested Tit vs Coal Tit | -0.10 ± 0.20 | -0.51 | 0.612 |
| Acidophilic broadleaves woods | Great Tit vs Marsh Tit | 0.09 ± 0.13 | 0.75 | 0.456 |
| Acidophilic broadleaves woods | Great Tit vs Blue Tit | 0.03 ± 0.15 | 0.20 | 0.844 |
| Acidophilic broadleaves woods | Great Tit vs Coal Tit | 0.13 ± 0.17 | 0.80 | 0.426 |

| | | | | |
|---|---|---|---|---|
| Acidophilic broadleaves woods | Great Tit vs Crested Tit | 0.19 ± 0.15 | 1.23 | 0.219 |
| Acidophilic broadleaves woods | Marsh Tit vs Blue Tit | -0.06 ± 0.16 | -0.39 | 0.695 |
| Acidophilic broadleaves woods | Marsh Tit vs Crested Tit | 0.09 ± 0.16 | 0.59 | 0.559 |
| Acidophilic broadleaves woods | Marsh Tit vs Coal Tit | 0.04 ± 0.18 | 0.23 | 0.819 |
| Acidophilic broadleaves woods | Blue Tit vs Crested Tit | 0.16 ± 0.18 | 0.86 | 0.390 |
| Acidophilic broadleaves woods | Blue Tit vs Coal Tit | 0.10 ± 0.20 | 0.53 | 0.598 |
| Acidophilic broadleaves woods | Crested Tit vs Coal Tit | -0.05 ± 0.20 | -0.28 | 0.783 |
| Mesophilic broadleaves woods | Great Tit vs Marsh Tit | -0.06 ± 0.13 | -0.48 | 0.633 |
| Mesophilic broadleaves woods | Great Tit vs Blue Tit | -0.23 ± 0.15 | -1.47 | 0.142 |
| Mesophilic broadleaves woods | Great Tit vs Coal Tit | 0.20 ± 0.17 | 1.17 | 0.242 |
| Mesophilic broadleaves woods | Great Tit vs Crested Tit | 0.25 ± 0.15 | 1.66 | 0.097 |
| Mesophilic broadleaves woods | Marsh Tit vs Blue Tit | -0.17 ± 0.16 | -1.02 | 0.308 |
| Mesophilic broadleaves woods | Marsh Tit vs Crested Tit | 0.31 ± 0.16 | 1.94 | 0.052 |
| Mesophilic broadleaves woods | Marsh Tit vs Coal Tit | 0.26 ± 0.18 | 1.46 | 0.145 |
| Mesophilic broadleaves woods | Blue Tit vs Crested Tit | 0.48 ± 0.18 | 2.60 | 0.009 |
| Mesophilic broadleaves woods | Blue Tit vs Coal Tit | 0.42 ± 0.20 | 2.14 | 0.032 |
| Mesophilic broadleaves woods | Crested Tit vs Coal Tit | -0.06 ± 0.20 | -0.29 | 0.772 |
| Alien species woods | Great Tit vs Marsh Tit | 0.13 ± 0.13 | 1.01 | 0.315 |
| Alien species woods | Great Tit vs Blue Tit | -0.06 ± 0.16 | -0.36 | 0.721 |
| Alien species woods | Great Tit vs Coal Tit | 0.48 ± 0.17 | 2.90 | 0.004 |
| Alien species woods | Great Tit vs Crested Tit | 0.49 ± 0.15 | 3.21 | 0.001 |
| Alien species woods | Marsh Tit vs Blue Tit | -0.06 ± 0.12 | -0.36 | 0.721 |
| Alien species woods | Marsh Tit vs Crested Tit | 0.36 ± 0.16 | 2.25 | 0.024 |
| Alien species woods | Marsh Tit vs Coal Tit | 0.36 ± 0.17 | 2.05 | 0.040 |
| Alien species woods | Blue Tit vs Crested Tit | 0.42 ± 0.18 | 2.30 | 0.022 |
| Alien species woods | Blue Tit vs Coal Tit | 0.42 ± 0.20 | 2.13 | 0.033 |
| Alien species woods | Crested Tit vs Coal Tit | 0.00 ± 0.20 | -0.02 | 0.987 |

**Table S3.** Percentage of territory overlap for each species pair within each territory mapping area. We calculated these proportions at species level, by merging territories of each species within each mapping area. Is noteworthy that the calculation of overlapping proportions was conducted asymmetrically, i.e. for each species, we determined the percentage of its territory covered by that of every other species. Note that Coal Tits were absent from area I and M, while Crested Tits were absent from area I.

**Area A**

| Vs | Great Tit | Marsh Tit | Crested Tit | Coal Tit | Blue Tit |
|---|---|---|---|---|---|
| **Great Tit** | - | 14.78 | 7.21 | 5.48 | 5.57 |
| **Marsh Tit** | 19.76 | - | 13.13 | 4.17 | 3.8 |
| **Crested Tit** | 16.03 | 21.83 | - | 4.29 | 0.60 |
| **Coal Tit** | 14.97 | 8.51 | 5.27 | - | 3.12 |
| **Blue Tit** | 22.05 | 11.25 | 1.06 | 4.52 | - |

**Area C**

| Vs | Great Tit | Marsh Tit | Crested Tit | Coal Tit | Blue Tit |
|---|---|---|---|---|---|
| **Great Tit** | - | 14.69 | 1.95 | 0 | 3.65 |
| **Marsh Tit** | 15.32 | - | 0.04 | 0 | 3.58 |
| **Crested Tit** | 2.57 | 0.05 | - | 0.73 | 0 |
| **Coal Tit** | 0 | 0 | 7.34 | - | 0 |
| **Blue Tit** | 11.48 | 10.8 | 0 | 0 | - |

**Area D**

| Vs | Great Tit | Marsh Tit | Crested Tit | Coal Tit | Blue Tit |
|---|---|---|---|---|---|
| **Great Tit** | - | 8.79 | 0 | 0.05 | 4.91 |
| **Marsh Tit** | 5.18 | - | 1.03 | 4.05 | 1.98 |
| **Crested Tit** | 0 | 5.1 | - | 0.14 | 0 |
| **Coal Tit** | 0.37 | 53.98 | 0.37 | - | 0 |
| **Blue Tit** | 8.93 | 6.12 | 0 | 0 | - |

**Area E**

| Vs | Great Tit | Marsh Tit | Crested Tit | Coal Tit | Blue Tit |
|---|---|---|---|---|---|
| **Great Tit** | - | 2.25 | 1.38 | 5.91 | 9.38 |
| **Marsh Tit** | 6.73 | - | 10.85 | 11.32 | 8.08 |
| **Crested Tit** | 1.29 | 3.38 | - | 5.59 | 4.29 |
| **Coal Tit** | 7.34 | 4.71 | 7.44 | - | 8.9 |
| **Blue Tit** | 23.66 | 6.82 | 11.6 | 18.07 | - |

**Area F**

| Vs | Great Tit | Marsh Tit | Crested Tit | Coal Tit | Blue Tit |
|---|---|---|---|---|---|
| **Great Tit** | - | 3.85 | 3.08 | 0 | 6.03 |
| **Marsh Tit** | 18.33 | - | 0 | 0 | 2.86 |
| **Crested Tit** | 8.87 | 0 | - | 5.03 | 2.39 |
| **Coal Tit** | 0 | 0 | 16.38 | - | 0 |
| **Blue Tit** | 17.5 | 1.74 | 2.41 | 0 | - |

**Area G**

| Vs | Great Tit | Marsh Tit | Crested Tit | Coal Tit | Blue Tit |
|---|---|---|---|---|---|
| **Great Tit** | - | 23.17 | 4.17 | 0 | 11.35 |
| **Marsh Tit** | 19.78 | - | 8.86 | 0 | 8.02 |
| **Crested Tit** | 3.76 | 9.36 | - | 0.4 | 1.15 |
| **Coal Tit** | 0 | 0 | 0.76 | - | 0.56 |
| **Blue Tit** | 15.06 | 12.47 | 1.69 | 0.43 | - |

**Area H**

| Vs | Great Tit | Marsh Tit | Crested Tit | Coal Tit | Blue Tit |
|---|---|---|---|---|---|
| **Great Tit** | - | 1.68 | 8.18 | 17.14 | 7.82 |
| **Marsh Tit** | 2.1 | - | 10.75 | 10.39 | 5.88 |
| **Crested Tit** | 4.37 | 4.58 | - | 3.89 | 13.57 |
| **Coal Tit** | 21.86 | 10.57 | 9.29 | - | 4.33 |
| **Blue Tit** | 14.12 | 8.48 | 45.87 | 6.13 | - |

**Area I**

| Vs | Great Tit | Marsh Tit | Crested Tit | Coal Tit | Blue Tit |
|---|---|---|---|---|---|
| **Great Tit** | - | 2.73 | / | / | 5.63 |
| **Marsh Tit** | 12.22 | - | / | / | 1.51 |
| **Crested Tit** | / | / | / | / | / |
| **Coal Tit** | / | / | / | / | / |
| **Blue Tit** | 25.27 | 1.51 | / | / | - |

**Area M**

| Vs | Great Tit | Marsh Tit | Crested Tit | Coal Tit | Blue Tit |
|---|---|---|---|---|---|
| **Great Tit** | - | 7.78 | 1.71 | / | 0.8 |
| **Marsh Tit** | 14.11 | - | 2.88 | / | 0 |
| **Crested Tit** | 26.55 | 24.67 | - | / | 0 |
| **Coal Tit** | / | / | / | / | / |
| **Blue Tit** | 3.96 | 0 | 0 | / | - |

**Area N**

| Vs | Great Tit | Marsh Tit | Crested Tit | Coal Tit | Blue Tit |
|---|---|---|---|---|---|
| **Great Tit** | - | 4.88 | 0 | 0 | 2.68 |
| **Marsh Tit** | 10.5 | - | 0.39 | 0 | 2.35 |
| **Crested Tit** | 0 | 1.26 | - | 1.4 | 0 |
| **Coal Tit** | 0 | 0 | 0.63 | - | 0 |
| **Blue Tit** | 16.88 | 6.88 | 0 | 0 | - |

**Appendix 1**

To test if % of overlap of breeding territories was associated with the density of territories across mapping areas we adopted the following approach.

Initially, we computed the following variables:

1. The average % of overlap for the "broadleaf" and "conifer species" per each area, calculated as the mean among the % overlaps for each species pair within its subgroup per each area.
2. Territory density for "broadleaf" and "conifer species", defined as the number of territories of species within the same subgroup on the extent (ha) of suitable habitat areas for each mapped area. Suitable habitat for "broadleaf species" included acidophilic mixed woods, acidophilic broadleaves woods, mesophilic broadleaves woods, and areas with alien plant species, while suitable habitat for "conifer species" included acidophilic conifer woods and acidophilic mixed woods. This means that for each area we could obtain a maximum of two territory density estimates.

Thus, we tested the correlation between the mean % of overlap and the territory density separately for "broadleaf" and "conifer species" by performing a Spearman correlation test. No significant effects were observed neither for "broadleaf" ($r_s$ = -0.01, n = 10, $P$ = 0.999) nor for "conifer species" ($r_s$ = 0.24, n = 8, $P$ = 0.582). All analyses were conducted using R version 4.2. (R Core Team 2023). Consequently, we can infer that the % of overlap was independent of the density of territories across mapping areas.

**Appendix 2**

In this section, we provide a detailed description of the methods employed to randomly relocate breeding territories of the species within each mapping area. All analyses were conducted using R version 4.2.0 (R Core Team 2023) with the "sf" package (Pebesma 2018). The purpose of this simulation was to estimate the % of territory overlap that would occur if species occupied territories independently and randomly of each other, thereby assuming the absence of spatial segregation of breeding territories. These estimated values were then compared within each area against the observed ones. The procedure involved the following steps.

1. Because the breeding territories presence, abundance and distribution strictly depend on the availability of suitable habitat, we first delineated the boundaries where to relocate territories based on potential suitable habitat types for each species within each mapping area, as determined by habitat use during the breeding period (see Results and Figure 1 in the Main Text, and Table S3). Specifically, we combined acidophilic mixed woods, acidophilic broadleaves woods, mesophilic broadleaves woods and alien plant species woods to define the relocation areas for "broadleaf species" (Great Tits, Blue Tits, and Marsh Tits) and acidophilic conifer woods and acidophilic mixed woods for "conifer species" (Crested Tits and Coal Tits). All open farmlands and urban patches were excluded. Hence, we identified specific forest portions for relocation within each subgroup (see Fig. S2). Notably, since Crested Tits and Coal Tits where found together only in 8 mapping areas out of 10, we relied only on these 8 sites.

2. A negative buffer, equivalent to the radius obtained from the average surface area of territories for each species in each area, was applied to prevent territory relocation outside the mapping area and unsuitable habitats.
3. Separately for each species within each area, we generated points corresponding to the number of observed territories, ensuring a minimum distance between points equivalent to twice the territory radius for that species, using the "spatstat" package (Baddeley *et al.* 2015). This approach aimed to simulate realistic territory spacing while allowing for some degree of overlap, consistent with observed territory behavior.
4. Subsequently, we randomly relocated observed territories by associating territory centroids with the generated points. This process involved shifting territories by calculating Euclidean distances between centroids and generated points, then adjusting the vertices of original territory polygons accordingly (see Fig. S3). We considered the possibility that territories, which vary in shape, might be connected to points on the edge of the relocation area, potentially extending beyond suitable habitat boundaries. However, this concern was alleviated by the observation of occasional inclusion of unsuitable habitats within observed territories.
5. At this point, we quantified the % of overlap between species within each subgroup. We first calculated the total surface area covered by breeding territories of each species and subsequently, we calculated the % of overlap for each species relative to other ones within each subgroup. Notably, the calculation of overlapping proportions was conducted asymmetrically, i.e. for each species, we determined the percentage of its territory covered by those of other species. For example, in area C, both Crested Tits and Coal Tits have the same number of territories each (N = 3), but Crested Tits occupied slightly larger territories. From Crested Tits' perspective, the overlap with Coal Tits overs a small portion of their territory (0.73 %), while

from the Coal Tits' perspective, the portion shared with Crested Tits is proportionally larger (7.34 %).

6. This relocation process was repeated 100 times for each species within each mapping area, resulting in 1,000 simulations for "broadleaf species" and 800 simulations for "conifer species" (see Fig. S4).
7. Finally, the average % overlap among relocated territories for each species pair comparison within each area was calculated and incorporated into the dataset along with observed overlaps.

These data were used to perform a series of Wilcoxon signed-rank tests comparing observed and simulated % overlap for each species pair comparison (separately for all "broadleaf" and "conifer species"). We preferred performing Wilcoxon test as it does not assume normality distribution for data. Tests were conducted for paired data in each mapping area (N = 10 for "broadleaf" and N = 8 for "conifer species"). Results are summarized in Table S5 and Fig. S5.

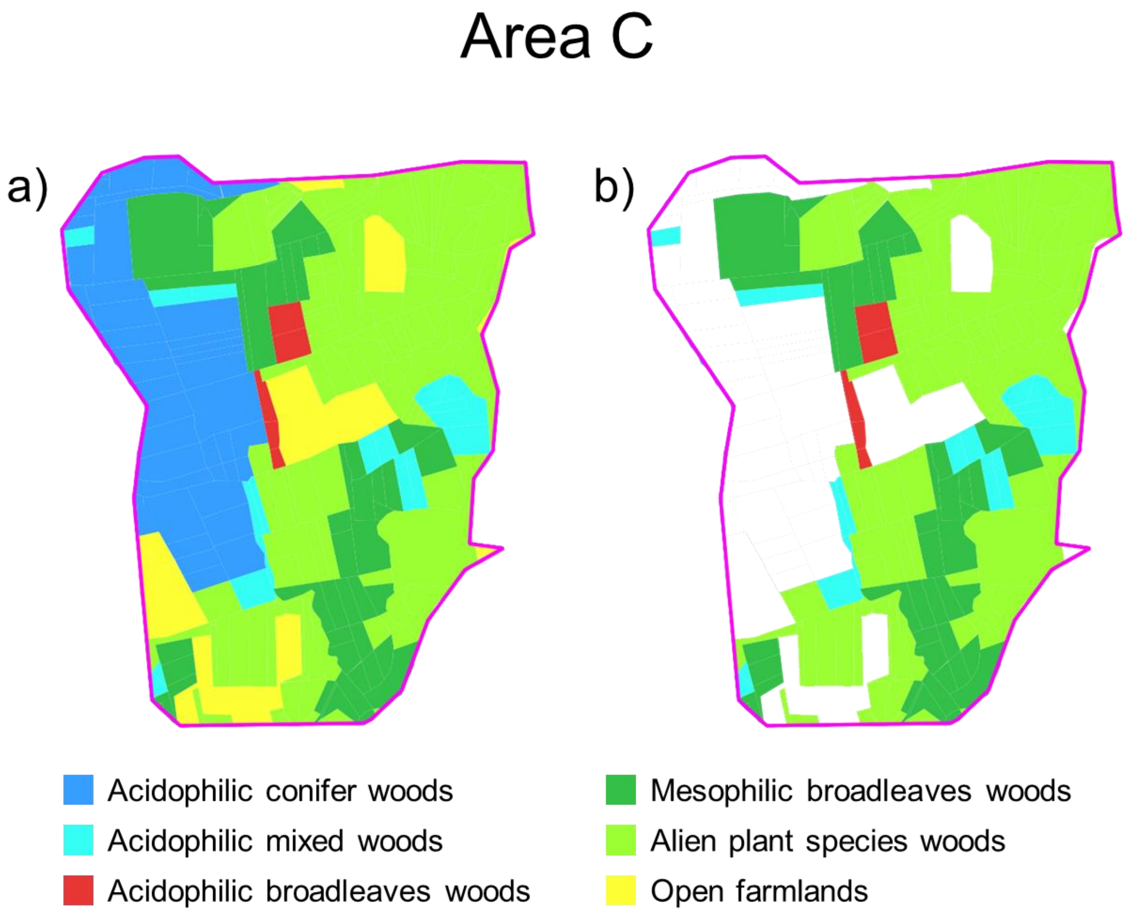


**Figure S2.** The figure shows (a) an example of a mapping area (area C) subdivided into the identified vegetation types, in which (b) suitable vegetation types were selected for the 3 species of the "broadleaf" subgroup (i.e. acidophilic mixed woods, acidophilic broadleaves woods, mesophilic broadleaves woods and alien plant species woods). This portion represents the area where the "broadleaf species" territories were relocated.

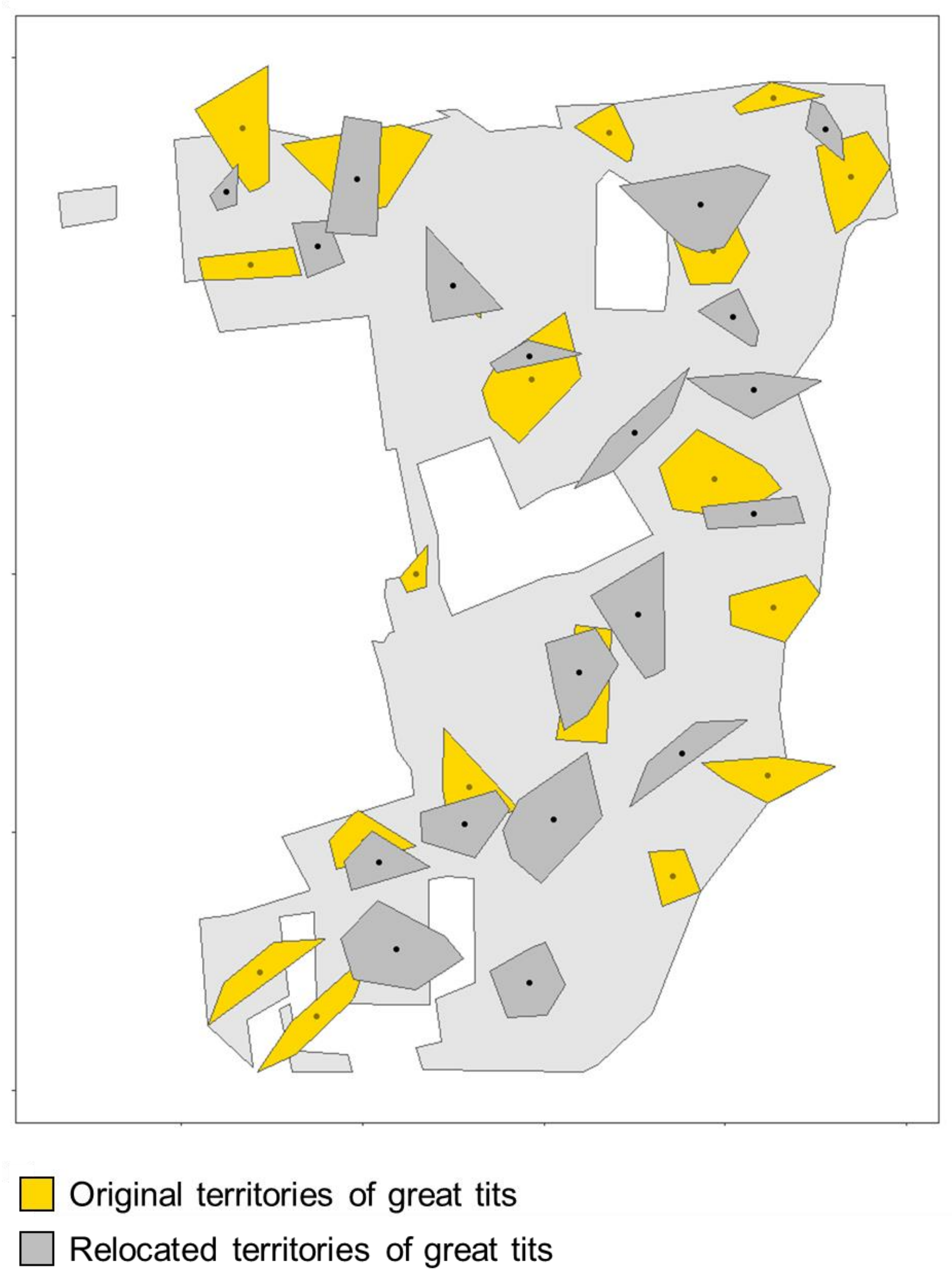


**Figure S3.** The figure shows the original tits territories together with those randomly relocated in a simulation for the mapping area C. Dark dots inside territories represent centroids.

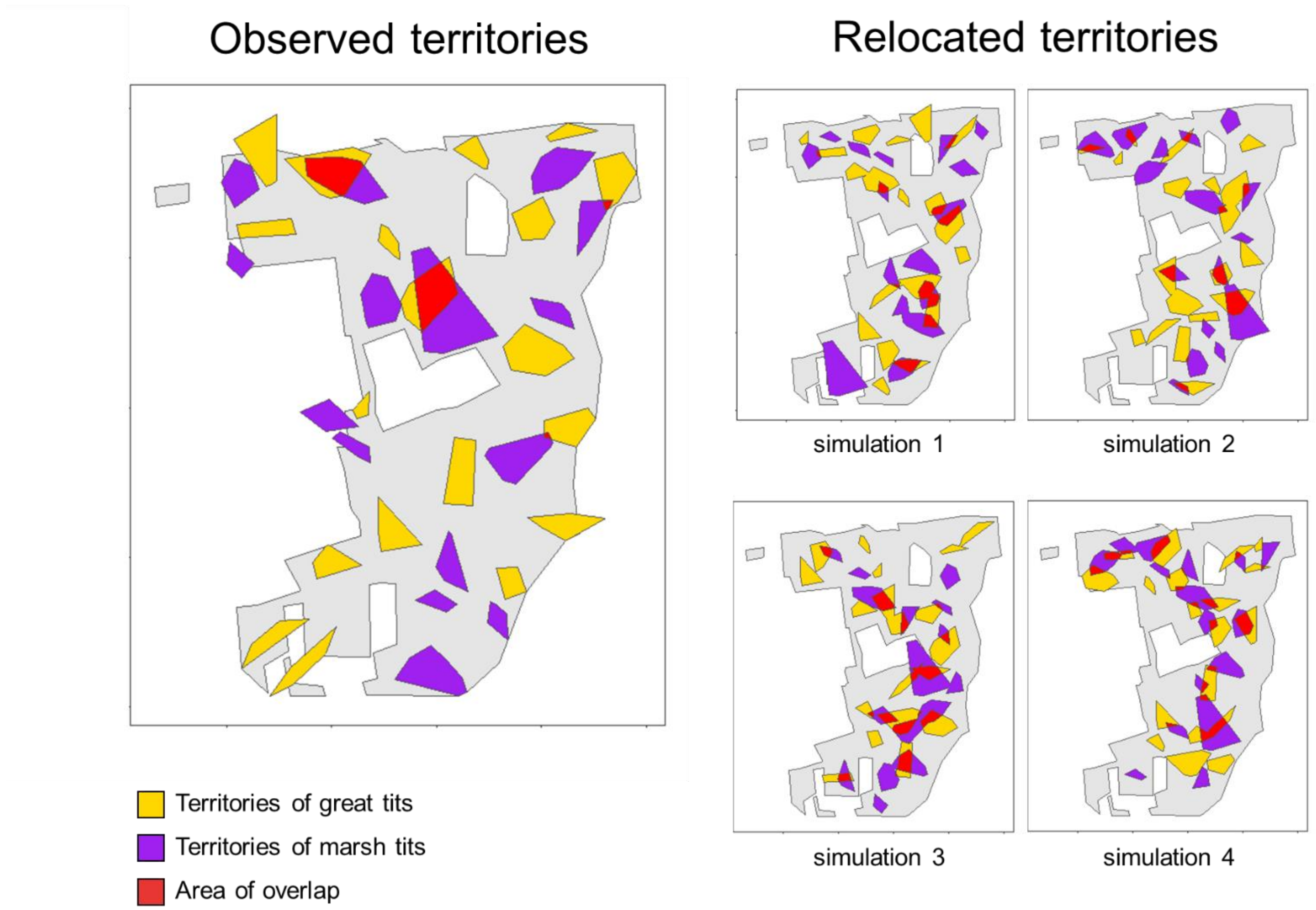


**Figure S4.** The figure on the left shows the distribution of the original Great Tits and Marsh Tit territories in the mapping area C, and their observed area of overlap. On the right-hand side, 4 maps are representing 4 examples of simulations in which territories of the two tit species were relocated randomly according to the procedure described in “Supporting methods”. Notably, the area of overlap for simulations is larger than the original.

**Table S4.** Summaries of Wilcoxon tests paired performed to fit the percentage of overlap calculated from observed territories with those from "simulated" data, for each comparison between species pairs. a) results of each species pairs comparison from the "broadleaf species" subgroup, b) results of "broadleaf species". Overlaps were expressed as the average percentage (± standard error) of a territory of a given species overlapping with a territory of other species within each territory mapping area. Observed overlaps resulted to be significantly ($P < 0.05$) lower than those obtained from "simulated" territories – see Appendix 2 and Fig. S5 for details.

| **a) "Broadleaf species"** | | | | |
|---|---|---|---|---|
| *Species pair comparison* | *Observed mean ± S.E.* | *Simulated mean ± S.E.* | *V* | *P* |
| Great Tit vs Blue Tit | 5.78 ± 0.99 | 8.13 ± 1.23 | 48 | 0.037 |
| Blue Tit vs Great Tit | 15.89 ± 2.11 | 23.04 ± 2.75 | 48 | 0.037 |
| Blue Tit vs Marsh Tit | 6.61 ± 1.38 | 15.87 ± 2.74 | 54 | 0.004 |
| Marsh Tit vs Blue Tit | 3.81 ± 0.86 | 8.62 ± 1.22 | 54 | 0.004 |
| Great Tit vs Marsh Tit | 8.46 ± 2.23 | 14.63 ± 2.52 | 55 | 0.002 |
| Marsh Tit vs Great Tit | 12.40 ± 1.97 | 22.51 ± 2.51 | 55 | 0.002 |
| **b) "Conifer species"** | | | | |
| *Species pair comparison* | *Observed mean ± S.E.* | *Simulated mean ± S.E.* | *V* | *P* |
| Coal Tit vs Crested Tit | 5.93 ± 1.94 | 15.27 ± 3.70 | 33 | 0.039 |
| Crested Tit vs Coal Tit | 2.68 ± 0.79 | 7.93 ± 1.87 | 33 | 0.039 |

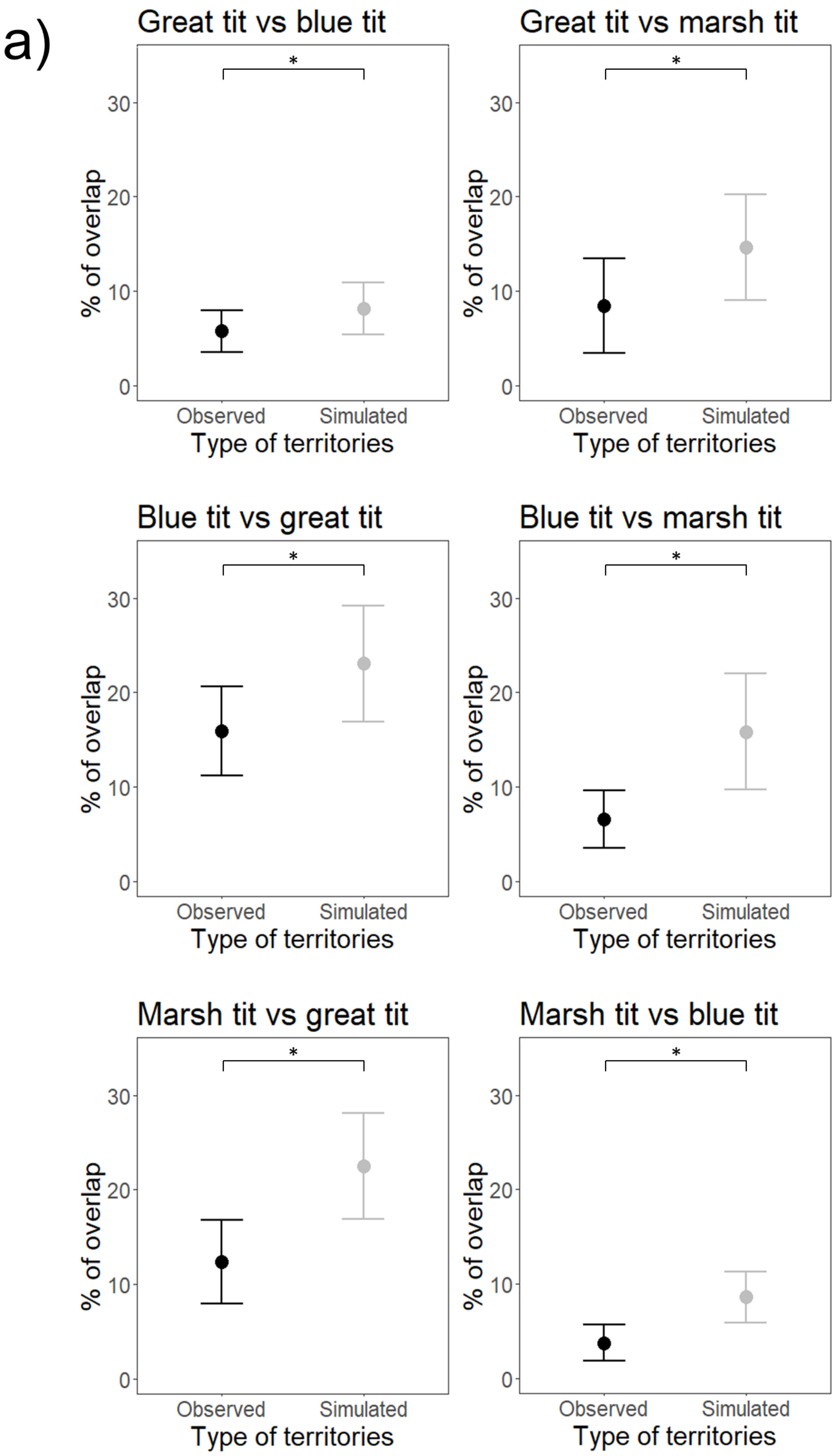
a)
Great tit vs blue tit
Great tit vs marsh tit
Blue tit vs great tit
Blue tit vs marsh tit
Marsh tit vs great tit
Marsh tit vs blue tit
*
% of overlap
0
10
20
30
Observed
Simulated
Type of territories

b)

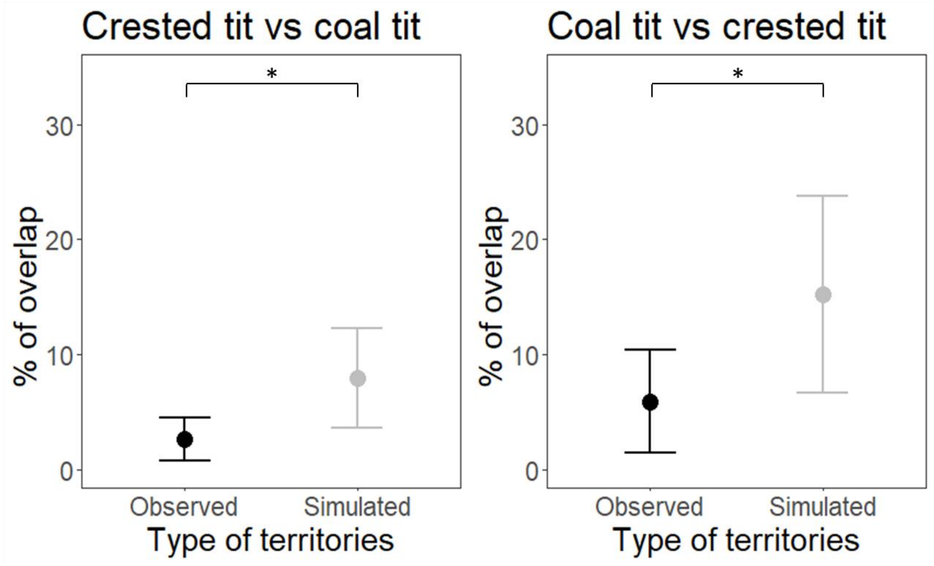


**Figure S5.** Comparisons between average values (with 95% CI) of % of overlap obtained from observed breeding territories (i.e. those from mapping procedure in each mapping areas of Figure S1) with those calculated from randomly relocated territories (see Supporting Methods), for each species pair. Species from the "broadleaf" subgroup (Great Tit, Blue Tit, Marsh Tit) were considered separately from the "conifer" species (Crested Tit, Coal Tit).

**Table S5.** Manly's selection ratios at the home-range scale during the non-breeding period according to species and habitat class. Locations = number of observations per each habitat class. See Table S1 for details on each habitat class.

| Species | Habitat class | Availability (ha) | Locations (N) | Selection ratio | Lower 95% CI | Upper 95% CI |
|---|---|---|---|---|---|---|
| Great Tit | Acidophilic conifer woods | 1249.37 | 44 | 1.237 | 0.941 | 1.601 |
| Great Tit | Acidophilic mixed woods | 680.02 | 34 | 1.756 | 1.278 | 2.375 |
| Great Tit | Acidophilic broadleaves woods | 534.23 | 41 | 2.695 | 2.026 | 3.530 |
| Great Tit | Mesophilic broadleaves woods | 436.44 | 21 | 1.690 | 1.119 | 2.513 |
| Great Tit | Alien species woods | 1072.07 | 56 | 1.834 | 1.448 | 2.288 |
| Blue Tit | Acidophilic conifer woods | 1249.37 | 7 | 0.323 | 0.158 | 0.644 |
| Blue Tit | Acidophilic mixed woods | 680.02 | 15 | 1.270 | 0.782 | 2.013 |
| Blue Tit | Acidophilic broadleaves woods | 534.23 | 33 | 3.557 | 2.606 | 4.735 |
| Blue Tit | Mesophilic broadleaves woods | 436.44 | 28 | 3.695 | 2.621 | 5.078 |
| Blue Tit | Alien species woods | 1072.07 | 11 | 0.591 | 0.334 | 1.020 |
| Marsh Tit | Acidophilic conifer woods | 1249.37 | 49 | 1.547 | 1.200 | 1.960 |
| Marsh Tit | Acidophilic mixed woods | 680.02 | 39 | 2.262 | 1.691 | 2.975 |
| Marsh Tit | Acidophilic broadleaves woods | 534.23 | 33 | 2.437 | 1.768 | 3.300 |
| Marsh Tit | Mesophilic broadleaves woods | 436.44 | 26 | 2.350 | 1.629 | 3.331 |
| Marsh Tit | Alien species woods | 1072.07 | 55 | 2.024 | 1.598 | 2.518 |
| Crested Tit | Acidophilic conifer woods | 1249.37 | 121 | 3.718 | 3.261 | 4.170 |
| Crested Tit | Acidophilic mixed woods | 680.02 | 56 | 2.833 | 2.239 | 3.525 |
| Crested Tit | Acidophilic broadleaves woods | 534.23 | 11 | 0.876 | 0.493 | 1.532 |
| Crested Tit | Mesophilic broadleaves woods | 436.44 | 13 | 1.001 | 0.590 | 1.670 |
| Crested Tit | Alien species woods | 1072.07 | 9 | 0.331 | 0.175 | 0.615 |
| Coal Tit | Acidophilic conifer woods | 1249.37 | 33 | 3.934 | 3.051 | 4.758 |
| Coal Tit | Acidophilic mixed woods | 680.02 | 2 | 0.438 | 0.121 | 1.490 |
| Coal Tit | Acidophilic broadleaves woods | 534.23 | 5 | 1.394 | 0.605 | 3.015 |
| Coal Tit | Mesophilic broadleaves woods | 436.44 | 6 | 2.048 | 0.956 | 4.115 |
| Coal Tit | Alien species woods | 1072.07 | 4 | 0.556 | 0.219 | 1.324 |

**Appendix 3**

Prior to conducting the analysis of the foraging habitat scale in "Habitat use and differentiation during the non-breeding period" (see Methods), a preliminary analysis was conducted to ensure that the distribution of tits in the canopy thirds during foraging did not differ significantly when individuals were observed alone versus in flocks. Within each species, we compared the distribution in the canopy portions used during foraging between observations of solitary individuals and those within flocks. The Cochran-Mantel-Haenzsel (CMH) test was employed to assess differences in canopy distribution. No differences were identified for the species during foraging in singletons or in flocks (Table S6).

**Table S6.** Differences in the distribution of individuals of different species foraging in tree canopy between observations considering flocks and singletons during the non-breeding period (Cochran-Mantel-Haenzsel test for 3-dimensional contingency tables; Mangiafico and Mangiafico, 2017). No statistical difference between in-flock and solo foraging are observed in winter.

| **Species** | $M^2$ | **df** | ***P*** |
|---|---|---|---|
| Great Tit | 0.83 | 2 | 0.660 |
| Blue Tit | 0.51 | 2 | 0.774 |
| Marsh Tit | 2.39 | 2 | 0.302 |
| Crested Tit | 3.70 | 2 | 0.157 |
| Coal Tit | 4.83 | 2 | 0.089 |

**R Core Team**. 2023. R: A language and environment for statistical computing. R Foundation for Statistical Computing, Vienna, Austria.
